# A Disinhibitory Circuit for Contextual Modulation in Primary Visual Cortex

**DOI:** 10.1101/2020.01.31.929166

**Authors:** Andreas J. Keller, Mario Dipoppa, Morgane M. Roth, Matthew S. Caudill, Alessandro Ingrosso, Kenneth D. Miller, Massimo Scanziani

## Abstract

Context guides perception by influencing the saliency of sensory stimuli. Accordingly, in visual cortex, responses to a stimulus are modulated by context, the visual scene surrounding the stimulus. Responses are suppressed when stimulus and surround are similar but not when they differ. The mechanisms that remove suppression when stimulus and surround differ remain unclear. Here we use optical recordings, manipulations, and computational modelling to show that a disinhibitory circuit consisting of vasoactive-intestinal-peptide-expressing (VIP) and somatostatin-expressing (SOM) inhibitory neurons modulates responses in mouse visual cortex depending on the similarity between stimulus and surround. When the stimulus and the surround are similar, VIP neurons are inactive and SOM neurons suppress excitatory neurons. However, when the stimulus and the surround differ, VIP neurons are active, thereby inhibiting SOM neurons and relieving excitatory neurons from suppression. We have identified a canonical cortical disinhibitory circuit which contributes to contextual modulation and may regulate perceptual saliency.

## Introduction

The perception of a sensory stimulus is markedly influenced by the context in which the stimulus is embedded. In the visual system, the context is the visual scene surrounding the stimulus. Through the influence of its surround-the same visual stimulus may be perceived as more or less salient, allowing it to pop out or merge with the rest of the visual scene (Figure 1A; Bergen and Julesz, 1983; Lamme, 1995; Treisman and Garry Gelade, 1980). This aspect of sensory processing represents a fundamental computation to extract meaning from visual scenes.

**Figure 1.**
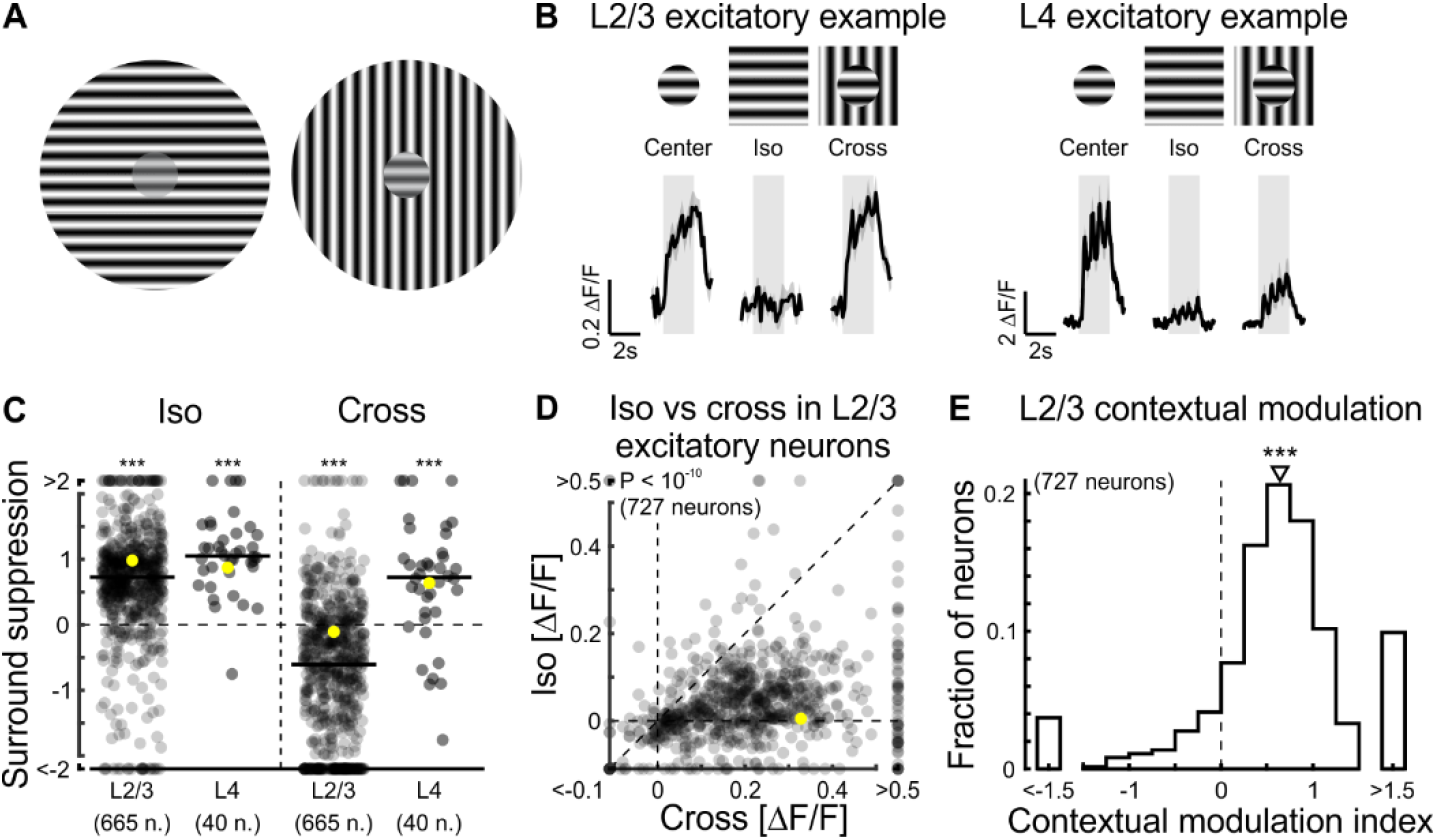
Contextual modulation in excitatory neurons. (A) The small grating patches in the centers had the same contrast but due to the distinct surround, they were perceived as more or less salient, allowing them to pop out (right) or merge with the rest of the visual scene (left). (B) Visual stimuli were presented to awake mice while imaging calcium responses in layer 2/3 (L2/3) excitatory neurons of primary visual cortex (V1) expressing GCaMP6f or GCaMP7f. Top: Schematic of a small grating patch (20° in diameter) presented alone (center), with an iso oriented surround (iso), or with a cross-oriented surround (cross). Bottom left: Trial-averaged calcium responses of an example L2/3 excitatory neuron to center, iso, and cross stimuli. Bottom right: Same but for an example layer 4 (L4) excitatory neuron. Here and in all other figures shaded areas are periods of stimulus presentation. (C) Surround suppression was computed for both L2/3 and L4 neurons as the difference in responses to center stimuli and the responses to iso (or cross) stimuli, normalized by the responses to center stimuli. Single-distribution two-sided Wilcoxon sign-rank test; iso L2/3, ***: p < 10^-10^; cross L2/3, ***: p < 10^-10^; 665 neurons in 9 mice; iso L4, ***: p < 10^-7^; cross L4, ***: p = 1.9 × 10^-4^; 40 neurons in 5 mice. Yellow symbols represent the example neurons shown in (B). Here and in all figures horizontal black lines indicate the median of the distribution. (D) Scatter plot of L2/3 responses to iso and cross. Paired two-sided Wilcoxon sign-rank test; p < 10^-10^ (727 neurons in 9 mice). Yellow symbol represents the example neuron shown in (B). (E) Contextual modulation index (CMI) was computed as the difference divided by the sum of the responses to cross and iso stimuli. Here and in all figures triangles above histograms indicate median. Single-distribution two-sided Wilcoxon sign-rank test; p < 10^-10^; same neurons as in (D).

Consistent with perceptual phenomena-neuronal responses to a visual stimulus are modulated by the visual scene surrounding the stimulus. This surround modulation occurs at several stages of the visual system including the retina (Alitto and Usrey, 2008; Chiao and Masland, 2003; Huang et al., 2019; McIlwain, 1964; Ölveczky et al., 2003; Solomon, 2006), the thalamus (Alitto and Usrey, 2008; Jones et al., 2012, 2015; Levick et al., 1972), and the visual cortex (Alexander and Van Leeuwen, 2010; Angelucci et al., 2017; Fitzpatrick, 2000; Kapadia et al., 2000; Knierim and van Essen, 1992; Rossi et al., 2001; Schnabel et al., 2018; Sillito et al., 1995), progressively increasing the complexity of the spatial features that are contextualized.

The classical feedforward receptive field (ffRF) of a neuron in primary visual cortex (V1) is the region in space in which a visual stimulus evokes a response (Hubel and Wiesel, 1962). The magnitude of this response can be modulated by stimulating the regions surrounding the ffRF. When a stimulus is large enough to cover both the ffRF and its surround, for example, the neuron’s responses are generally suppressed. This phenomenon, called surround suppression, is a well-established example of surround modulation (Blakemore and Tobin, 1972; Hubel and Wiesel, 1965; Kapadia et al., 1999; Knierim and van Essen, 1992; Nelson and Frost, 1978). It has been shown that anatomical substrates for surround suppression include feedback connections (Angelucci et al., 2017; Keller et al., 2020; Nurminen et al., 2018; Vangeneugden et al., 2019), interlaminar connections (Bolz and Gilbert, 1986) and specific subtypes of inhibitory neurons (Adesnik et al., 2012; Haider et al., 2010). The tuning properties of SOM inhibitory neurons (Adesnik et al., 2012; Dipoppa et al., 2018; Keller et al., 2020) and the fact that they connect to nearly all nearby excitatory neurons (Fino et al., 2013) make them ideal to contribute to surround suppression. Indeed, functional elimination of SOM neurons partially relieves excitatory neurons from surround suppression (Adesnik et al., 2012).

However, not all combinations of stimuli in the ffRF and surround generate suppression. Surround suppression occurs when the stimulus in the ffRF and in the surround share similar features. For example, the response of a neuron to a grating stimulus of a given orientation in its ffRF is suppressed when stimulating the surround with a grating of similar orientation. When the orientation of the grating in the surround differs from that in the ffRF, the response of the neuron is much less or no longer suppressed (Self et al., 2014; Sillito et al., 1995; Walker et al., 1999). Thus, the magnitude of surround suppression depends on the visual scene surrounding the stimulus in the ffRF. The mechanism that regulates surround suppression depending on the similarity between the stimulus in the ffRF and that in the surround remains elusive. We refer to this phenomenon as “contextual modulation”. To investigate the mechanisms of contextual modulation, we presented visual stimuli with different surrounds to awake mice while imaging calcium responses in excitatory and inhibitory neurons of V1. We focused on the three major classes of inhibitory neurons, parvalbumin-expressing (PV), SOM and VIP neurons (Lee et al., 2010; Pfeffer et al., 2013). PV and SOM neurons are the two principal sources of inhibition of cortical excitatory neurons in mouse V1. In contrast, VIP neurons primarily provide inhibition to SOM neurons, thus representing a key component of cortical disinhibitory circuits (Karnani et al., 2016; Pfeffer et al., 2013; Pi et al., 2013). We show that the responses of VIP and PV neurons were only suppressed by surrounds that shared similar features to the stimulus presented in the ffRF but not when they differed, as observed in excitatory neurons. Strikingly, the responses of SOM neurons were modulated in a manner opposite to all other neuron types, being specifically suppressed by surrounding stimuli that differ from those in the ffRF. To determine whether the interaction between VIP and SOM neurons could account for the contextual modulation observed in excitatory neurons, we developed a circuit model respecting biological constraints, which we trained to reproduce our measurements. Our model predicted that silencing VIP neurons would reduce contextual modulation in excitatory neurons. Consistent with this model, when VIP neurons were silenced optogenetically in V1, surround suppression in excitatory neurons became less sensitive to the stimulus features in the surround, thereby reducing contextual modulation. Thus, we show that a canonical cortical disinhibitory circuit critically contributes to the contextual modulation of excitatory neurons in V1.

## Results

### Contextual modulation in excitatory neurons

To assess contextual modulation in V1, we recorded from layer 2/3 (L2/3) excitatory neurons in awake head-fixed mice with two-photon calcium imaging. Contextual modulation was assessed by comparing the responses of individual neurons to small patches of oriented gratings (20° in diameter) presented alone (“center stimulus”), or together with two different surrounds: An iso-oriented surround (“iso stimulus”; i.e. a grating in the surround whose orientation matches that of the grating in the center), or a cross-oriented surround (“cross stimulus”; i.e. a grating in the surround whose orientation is orthogonal relative to that of the grating in the center; Figure 1B, top). The location of the center stimulus was centered on the ffRFs of the neurons (see Methods). The magnitude of the response of L2/3 excitatory neurons to center stimuli alone was larger than that to iso stimuli, consistent with iso stimuli generating surround suppression (Figure 1B, left). In contrast, the response to cross stimuli was similar to the response to center stimuli alone, consistent with the fact that cross stimuli generate less or no surround suppression than iso-stimuli, as previously described (Self et al., 2014; Sillito et al., 1995; Walker et al., 1999). We computed the magnitude of surround suppression as the difference in response to center stimuli and response to iso or cross stimuli, normalized by the response to center stimuli. Accordingly, surround suppression in L2/3 excitatory neurons was larger for iso stimuli than for cross stimuli (Figure 1C, D). To compare the modulation by the iso surround to that of the cross surround, we defined a contextual modulation index (CMI) for each neuron (Figure 1E; see Methods). The distribution of CMIs of excitatory neurons was skewed to positive values, indicating that their responses were stronger to the cross than to the iso stimulus. Since the distribution of CMIs was similar irrespective of whether or not the orientation of the center stimulus matched the neuron’s orientation preference (Figure S1), our analysis includes neurons independently of their orientation preference. Overall, excitatory neurons in L2/3 were strongly modulated by context, i.e. the strength of their responses depended on the features of the surround relative to those in the center.

To what extent is the contextual modulation of excitatory L2/3 neurons inherited from earlier stages of cortical processing? To answer this question, we measured the responses of excitatory neurons in layer 4 (L4), the main thalamic input layer, to center, iso and cross stimuli (Figure 1B, right). While L2/3 neurons, on average, were only suppressed by the iso stimulus, L4 neurons showed suppression in response to both iso and cross stimuli (Figure 1C). Thus, contextual modulation of L2/3 neurons is unlikely to be entirely inherited from L4 and may rely on local circuitry.

### Complementary contextual modulation in SOM and VIP neurons

What relieves L2/3 excitatory neurons from surround suppression when the stimulus in the surround differs from the stimulus in the center? Since surround suppression of L2/3 excitatory neurons relies, at least in part, on the activation of SOM inhibitory neurons (Adesnik et al., 2012), we compared the response of SOM neurons to iso and cross stimuli. We thus repeated the visual stimulation protocol used above while recording in SOM neurons (Figure 2A-C). Strikingly, the responses of SOM neurons to iso and cross stimuli were opposite to what we observed in excitatory neurons. While iso stimuli elicited strong responses in SOM neurons, as previously observed (Adesnik et al., 2012; Dipoppa et al., 2018; Keller et al., 2020), cross stimuli elicited smaller responses (Figure 2B). Accordingly, the distribution of their CMIs was shifted towards negative values (Figure 2C). The smaller response of SOM neurons to cross than to iso stimuli was not a general characteristic of inhibitory neurons. PV neurons, the other large class of inhibitory neurons that targets excitatory neurons in mouse V1 (Pfeffer et al., 2013), showed larger responses to cross than to iso stimuli (Figure 2D, E). Therefore, the distribution of their CMIs was shifted towards positive values (Figure 2F), similar to excitatory neurons and opposite to SOM neurons. Thus, SOM neurons are unique in the way they respond to cross and iso stimuli. What prevents SOM neurons from responding to cross as much as to iso stimuli? SOM neurons receive excitatory input from L2/3 neurons. Given that L2/3 neurons strongly respond to cross stimuli, it is unlikely that the excitatory input to SOM neurons is reduced in response to cross stimuli. We thus reasoned that cross stimuli may generate inhibition onto SOM neurons. VIP inhibitory neurons are a class of cortical neurons that preferentially inhibits other inhibitory neurons, including SOM neurons (Pfeffer et al., 2013). If VIP neurons prevent SOM neurons from responding to cross but not to iso stimuli, they should be more excited by cross than by iso stimuli. To test this hypothesis, we repeated the visual stimulation protocol used above while recording in VIP neurons. Consistent with our prediction, VIP neurons responded more strongly to cross than to iso stimuli, as shown by their positively shifted CMI (Figure 2G-I).

**Figure 2.**
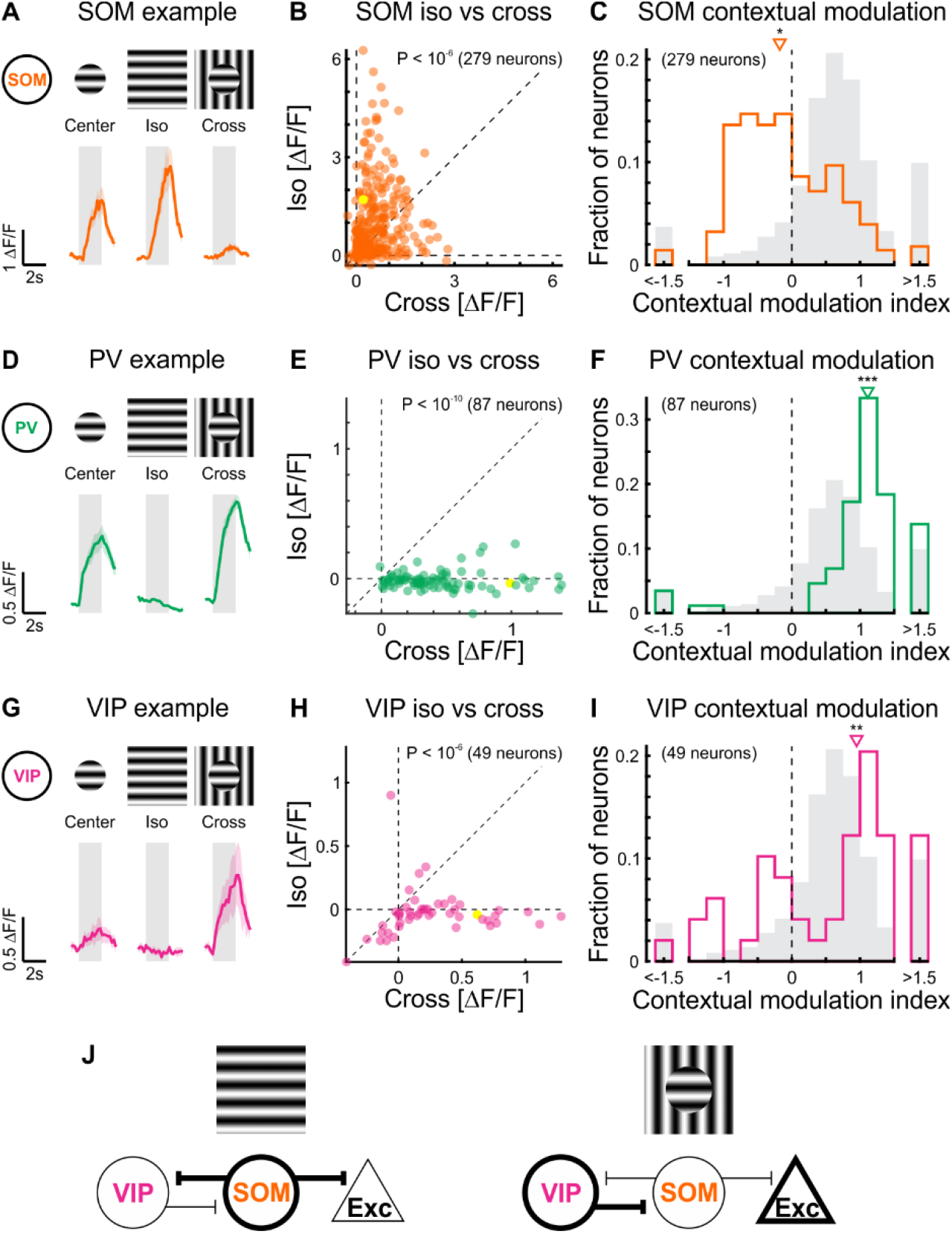
Contextual modulation in inhibitory neurons. (A) Top: Schematic of visual stimuli. Bottom: Trial-averaged calcium responses of an example somatostatin-expressing (SOM) inhibitory neuron expressing GCaMP6f to center, iso, and cross stimuli. (B) Scatter plot of the responses to iso and cross stimuli. Paired two-sided Wilcoxon signrank test; p < 10^-6^; 279 neurons in 13 mice. Yellow symbol represents the example neuron shown in (A). (C) CMI distribution of SOM neurons. Single-distribution two-sided Wilcoxon sign-rank test; *: p = 0.0081; same neurons as in (B). Gray shading: CMI distribution of L2/3 excitatory neurons (Figure 1E). (D-F) As above, but for parvalbumin-expressing (PV) inhibitory neurons. (E) Paired two-sided Wilcoxon sign-rank test; p – 10^-10^; 87 neurons in 9 mice. (F) Single-distribution two-sided Wilcoxon sign-rank test; ***: p < 10^-10^; same neurons as in (E). (G-I) As above, but for vasoactive-intestinal-peptide-expressing (VIP) inhibitory neurons. (H) Paired two-sided Wilcoxon sign-rank test; p < 10^-6^; 49 neurons in 6 mice. (I) Single-distribution two-sided Wilcoxon signrank test; **: p = 0.0012; same neurons as in (H). (J) Proposed mechanism of contextual modulation of excitatory neurons through the interaction between VIP and SOM neurons. Left: In response to an iso stimulus, SOM neurons are strongly driven and inhibit both VIP and excitatory neurons. Right: In response to the cross stimulus, VIP neurons are strongly driven and inhibit SOM neurons. Suppression of SOM neurons in turn disinhibits excitatory neurons

Taken together, these results are consistent with a mechanism in which the response modulation by the visual stimulus surrounding the ffRF of excitatory neurons is controlled by a disinhibitory circuit. The strong activation of SOM neurons by iso stimuli inhibits excitatory neurons thereby contributing to surround suppression (Figure 2J, left). In contrast, the strong activation of VIP neurons by cross stimuli inhibits SOM neurons, leading to the disinhibition of excitatory neurons (Figure 2J, right). A central prediction of this mechanism is that removing inhibition onto SOM neurons by functionally eliminating VIP neurons should lead to the suppression of excitatory neurons not only in response to iso stimuli but also in response to cross stimuli. In other words, functionally eliminating VIP neurons should reduce the response of excitatory neurons to cross stimuli more than that to iso stimuli.

### A circuit model predicts a role of VIP in contextual modulation

To test our intuition that the VIP-SOM disinhibitory circuit contributes to contextual modulation in L2/3 excitatory neurons, we developed a circuit model in which the model ‘units’ had supralinear input-output functions, consistent with experimental results (Adesnik, 2017; Priebe and Ferster, 2008; Priebe et al., 2004). Each unit of the circuit represented the average activity of a given neuron type (i.e. L2/3 excitatory, VIP, SOM, and PV neurons and L4 excitatory neurons), integrated in a ‘subnetwork’ with the other unit types (Figure 3A). Four such subnetworks were each assigned to one of two spatial locations (each considered the ‘surround’ of the other) and one of two preferred orientations (that were orthogonal to each other; Figure 3B). For the units sharing the same spatial location (both within and across subnetworks), we allowed all connections except those known to be weak (Adesnik et al., 2012; Karnani et al., 2016; Pfeffer et al., 2013). Subnetworks across spatial locations were connected only through L2/3 excitatory projections. We optimized the synaptic strengths between model units to match their responses to those observed experimentally. To determine the optimal synaptic strengths, we used a two-step procedure. We first generated many candidate solutions by performing non-negative regression (non-negative least squares), similarly to a previous study (Dipoppa et al., 2018), but on many sets of pseudo data obtained by randomly perturbing the experimental data. We then used the best solutions as initial conditions for a gradient-based optimization in a recurrent neural network (RNN; backpropagation through time with convolutional connections; Spoerer et al., 2017; see Methods). The top 15 models with the closest fits to the experimental data were used for further analysis (Figures 3C, S2A). These models had strong recurrent excitatory connections within a subnetwork (Figures 3D, S2B), consistent with previous studies (Cossell et al., 2015; Hofer et al., 2011; Ko et al., 2011; Peron et al., 2020). These strong recurrent connections led the circuit to become an inhibition-stabilized network for almost all top solutions, as has been found to underlie surround suppression (Adesnik, 2017; Ozeki et al., 2009). The combination of the supralinear input-output function and the inhibition stabilization mean that the circuit is a supralinear stabilized network (Ahmadian et al., 2013; Rubin et al., 2015).

**Figure 3.**
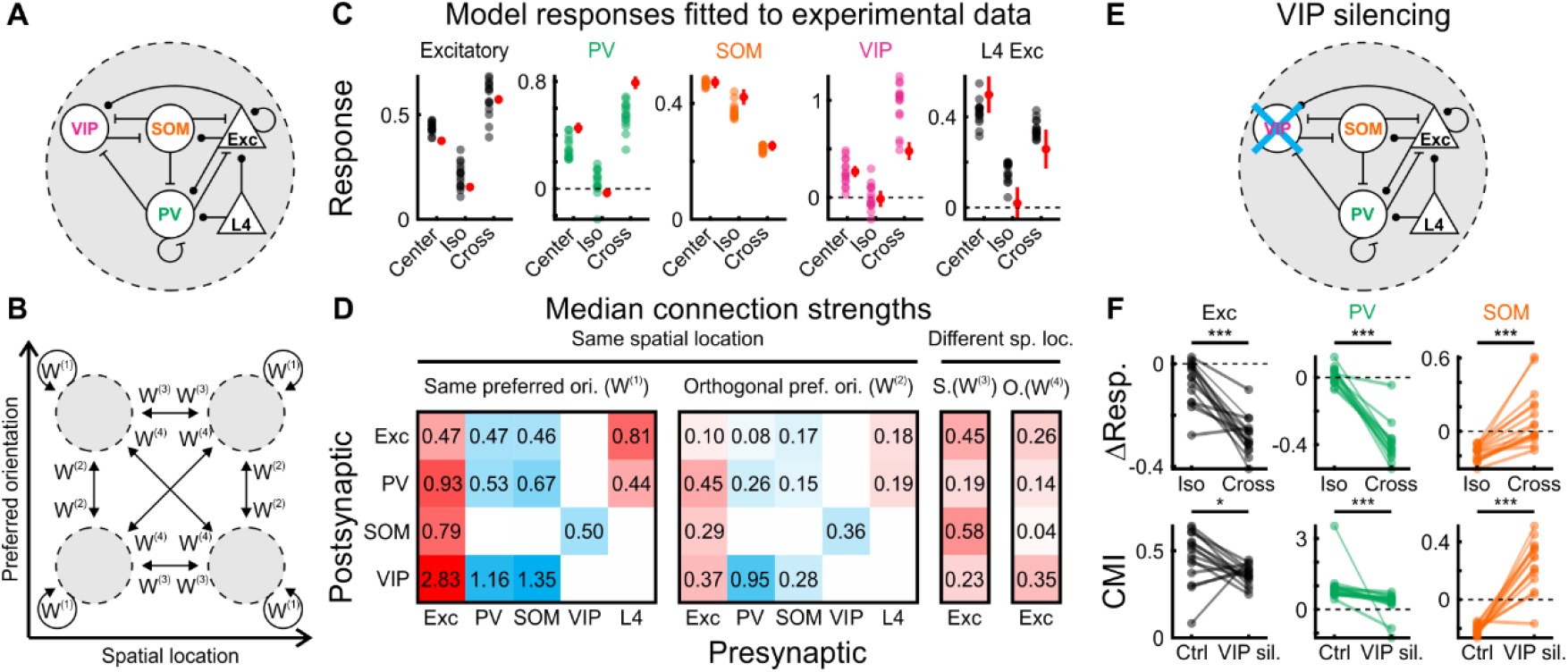
A computational model trained to fit experimental data predicts a role of VIP neurons in contextual modulation. (A) ‘Subnetwork’ of the model. Five unit types, L2/3 and L4 excitatory, VIP, SOM, and PV inhibitory units form a subnetwork. Unit types were connected according to biological constraints. (B) Four subnetworks were assigned to one of two spatial locations of the ffRF and one of two preferred orientations, connected with the weight matrices W^(1)^, W^(2)^, W^(3)^, and W^(4)^. (C) Responses of the different unit types from the top 15 models when the center, iso, and cross stimuli were centered on their spatial location and presented at their preferred orientation (see Figure S2A for the responses of all 4 subnetworks). Each dot represents the response of a unit from a single model. Red symbols represent experimental data (mean ± SEM; 317 excitatory neurons in 9 mice, 48 PV neurons in 9 mice, 200 SOM neurons in 13 mice, 30 VIP neurons in 6 mice, 22 L4 excitatory neurons in 5 mice). (D) Median connection strengths of the best 15 models. Excitatory connections are represented in red, inhibitory connections in blue. The 4 matrices correspond to W^(1)^, W^(2)^, W^(3)^, and W^(4)^ in (B). Note that this is not a working solution *per se* but the median connection strengths of the top 15 solutions. For an example solution see Figure S2B. (E) VIP units were silenced by effectively removing them from the circuit. (F) Top: Changes in response to iso and cross stimuli upon VIP silencing of L2/3 excitatory, PV and SOM unit types for the best 15 models. Bottom: CMI under control conditions compared to CMI during VIP silencing for the same unit types.

To determine the role of VIP units in the contextual modulation of excitatory units, we set the activity of VIP units to zero (Figure 3E). Silencing VIP units caused a larger absolute decrease in responses of excitatory units to cross than to iso stimuli (Figures 3F, top, S3A). While PV units were affected similarly to excitatory units, SOM units showed the opposite changes. In principle, a stronger absolute reduction in responses of excitatory neurons to cross than to iso stimuli is consistent with two possibilities. VIP units could simply regulate the overall gain in the network, that is, having the same relative impact on the responses of excitatory neurons to cross and iso stimuli. Alternatively, they could differentially regulate the responses of excitatory neurons depending on the stimulus. To distinguish between these two possibilities, we compared the CMIs of the different units under control conditions with the CMIs during VIP silencing. Consistent with VIP units differentially regulating the responses to iso and cross stimuli, their silencing decreased the CMI of excitatory units. This indicates that cross responses decreased proportionately more than iso responses. While PV units showed a decrease similar to excitatory units, the CMI of SOM units increased (Figure 3F, bottom).

The reduction of CMI in excitatory units upon VIP silencing was a prominent feature of the top 15 solutions but not of the next 85. While the majority of those 85 solutions showed a greater absolute reduction in responses of excitatory units to cross than to iso stimuli (Figure S3A), most of them did not show a decrease in CMI, but instead showed an increase (Figure S3B). By definition, those 85 solutions had a larger error in fitting the neuronal responses to visual stimuli than the top 15 solutions (Figure S3C). Thus, an important test of the models that best fit the data is whether silencing of VIP neurons in mouse V1 decreases the CMI of excitatory neurons.

By computing the sparse regression of these differences in CMI in excitatory units across these top 100 solutions against their synaptic strengths, we found that the changes in CMI could be well predicted by the values of a specific set of connection weights (Figure S3D, E). A reduction in CMI in excitatory units was correlated, for example, with a strengthening of VIP to SOM connections (Figure S3E). Indeed, such strengthening reduced CMI across these top 100 models (Figure S3F), indicating the importance of the VIP-SOM disinhibitory circuit in contextual modulation. Taken together, these results predict that silencing of VIP neurons in mouse V1 reduce contextual modulation of excitatory neurons.

### Inhibition of SOM neurons by VIP neurons contributes to contextual modulation

Does the functional elimination of VIP neurons preferentially decrease the response of excitatory neurons to cross stimuli compared to iso stimuli as predicted by the model? Since excitatory neurons are already almost maximally suppressed by iso stimuli, we reduced the contrast of all stimuli to 50%. This reduced the suppression of excitatory neurons by iso stimuli (suppression with iso stimuli; 100% contrast: 0.85 ± 0.02; 50% contrast: 0.58 ± 0.08; mean ± SEM; paired two-sided Wilcoxon sign-rank; p < 10^-10^; 641 neurons in 6 mice), consistent with previous observations (Kapadia et al., 1999), and allowed us to better compare the impact of silencing VIP neurons on the response to iso and cross stimuli. We optogenetically suppressed VIP neurons while recording their activity and the activity of putative excitatory neurons (Figure 4A). To determine the efficiency of optogenetic silencing of VIP neurons, we recorded their responses to center, iso and cross stimuli with and without photoactivation of the inhibitory opsin (see Methods). Photoactivation reduced both baseline activity as well as stimulus evoked responses of VIP neurons (Figure S4A-D). Furthermore, consistent with a previous study (Attinger et al., 2017), silencing VIP neurons had a suppressive effect on the baseline activity of putative excitatory neurons, confirming the disinhibitory impact of VIP neurons (Figure S4E, H). Strikingly, and consistent with the predictions of our model, silencing VIP neurons reduced the responses of putative excitatory neurons to cross stimuli significantly more than those to iso stimuli (Figure 4B, C). Importantly, as in our model, silencing VIP neurons also reduced the CMI of excitatory neurons, indicating that VIP neurons regulate the network in a context dependent manner (Figure 4D, E; also true for 100% contrast, Figure S4E-G). During this manipulation, excitatory neurons shifted their CMI towards zero (Figure 4E), implying that their responses were less dependent on the specific features of the surround.

**Figure 4.**
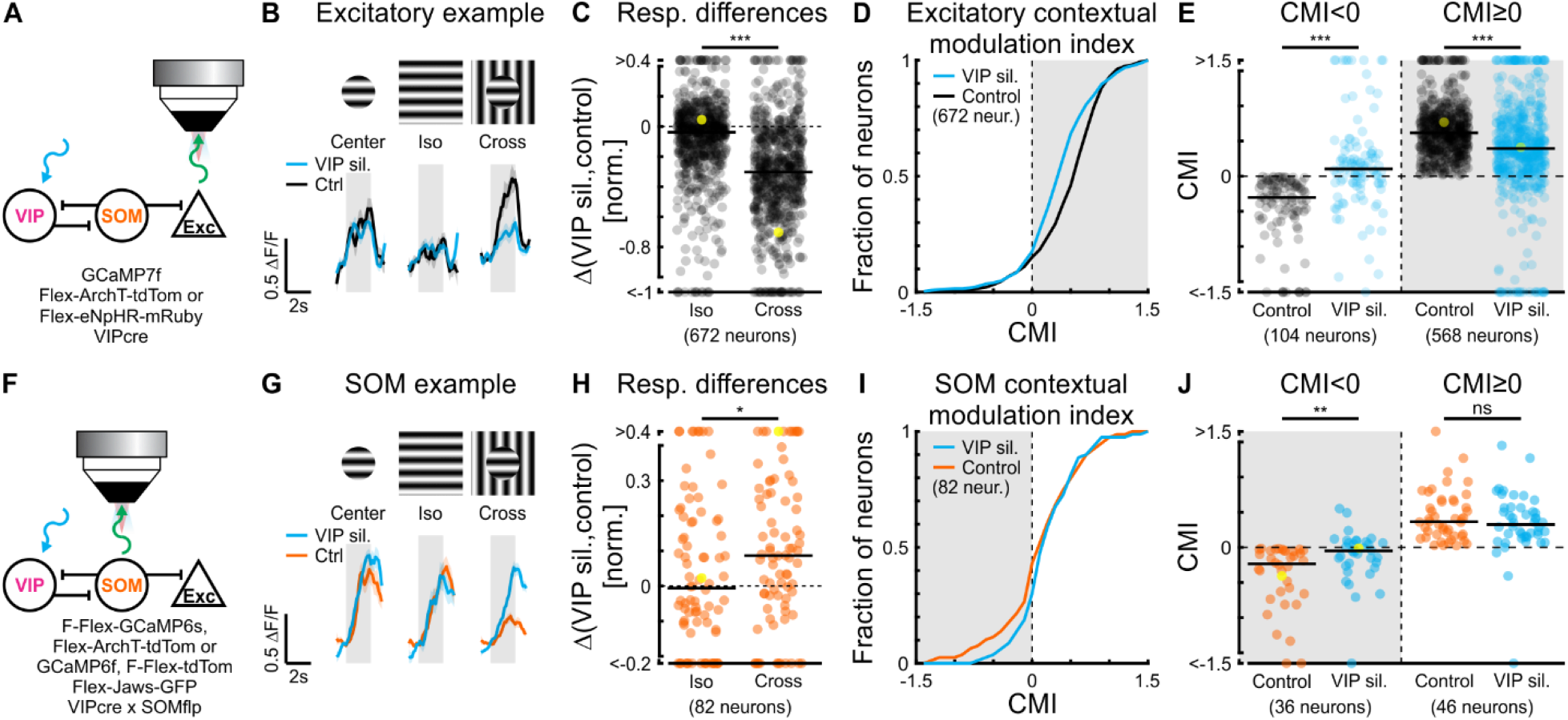
VIP and SOM neurons cooperatively contribute to contextual modulation in excitatory neurons. (A) Experimental setup. We conditionally expressed an inhibitory opsin, ArchT or eNpHR, in VIP neurons and unconditionally expressed a calcium indicator, GCaMP7f. (B) Trial-averaged calcium responses of a putative L2/3 excitatory neuron with and without silencing VIP neurons. Here, stimuli were presented at 50% contrast (similar responses to 100% stimuli, Figure S4E-G). (C) Iso and cross response differences between silencing VIP neurons and control conditions for putative excitatory neurons. Paired two-sided Wilcoxon sign-rank test; ***: p < 10^-10^; 672 neurons in 6 mice. Yellow symbol represents the example neuron shown in (B). (D) Cumulative sum of CMI in putative excitatory neurons. Same neurons as in (C). (E) Upon silencing VIP neurons, putative L2/3 excitatory neurons with a negative CMI increased their CMI and those with positive CMI decreased their CMI (gray shading). Paired two-sided Wilcoxon sign-rank; CMI<0 and CMI≥0, ***: p < 10^-10^; 104 and 568 neurons, respectively, in 6 mice. Yellow symbol represents the example neuron shown in (B). (F) Experimental setup. We conditionally expressed an inhibitory opsin, Jaws, in VIP neurons, conditionally expressed a red fluorescent reporter, tdTomato, in SOM neurons, and unconditionally expressed a calcium indicator, GCaMP6f; or we conditionally expressed an inhibitory opsin, ArchT, in VIP and conditionally expressed a calcium indicator, GCaMP6s, in SOM neurons. (G-J) Same as (B-D), but for SOM neurons. (H) Paired two-sided Wilcoxon sign-rank test; *: p = 0.027; 82 neurons in 8 mice. Yellow symbol represents the example neuron shown in (G). (J) Paired two-sided Wilcoxon sign-rank test; CMI<0, **: p = 0.0016; 36 neurons in 6 mice; CMI≥0, ns: p = 0.27; 46 neurons in 8 mice. Yellow symbol represents the example neuron shown in (G).

To determine whether the perturbation of VIP activity affects the activity of SOM neurons, we repeated our silencing protocol, however, this time, while recording from SOM neurons (Figure 4F). Upon VIP silencing, SOM neurons were significantly less suppressed by cross stimuli than by iso stimuli (Figure 4G, H). Moreover, SOM neurons with a negative CMI, which dominated the overall sample of SOM neurons (Figure 2C), shifted their CMI towards zero, while the ones with positive CMIs did not change on average (Figure 4I, J). Thus, the preferential suppression of SOM neurons by cross stimuli relies, at least in part, on the preferential activation of VIP neurons by these stimuli.

Taken together, based on optogenetic perturbations and computational modelling, these results demonstrate that the VIP-SOM disinhibitory circuit contributes to contextual modulation in excitatory neurons.

## Discussion

This study provides a mechanism for contextual modulation in V1 and reveals a disinhibitory circuit as a key mediator. Using imaging, optogenetic manipulations, and computational modeling, we find that the relationship between VIP and SOM inhibitory neurons contributes to the response profiles of L2/3 excitatory neurons in V1. When a uniform full-field stimulus is presented, VIP neurons are silent, while SOM neurons dominate the network and inhibit excitatory neurons. With a discontinuity in orientation between center and surround, VIP neurons are excited, inhibiting SOM neurons and effectively relieving excitatory neurons from SOM inhibition.

### Local circuits

The connectivity motifs between inhibitory neurons has been previously described (Jiang et al., 2015; Pfeffer et al., 2013; Pi et al., 2013) and are consistent with our findings. SOM neurons inhibit all other classes of neurons in L2/3 while VIP neurons preferentially inhibit SOM neurons. In addition, SOM neurons receive excitatory input from L2/3 neurons distributed over a relatively large retinotopic space (Adesnik et al., 2012). Our study indicates that when SOM neurons prevail over VIP neurons, excitatory neurons are inhibited, i.e. surround suppressed. Conversely, when VIP neurons prevail over SOM neurons, excitatory neurons are relieved from suppression.

What tips the balance in favor of VIP rather than SOM neurons in response to cross stimuli? Excitatory neurons in L4 are suppressed by cross stimuli, whereas L2/3 excitatory neurons are most active during cross compared to the other stimuli. In our model, excitatory drive to L2/3 excitatory neurons originating from L4 and from different spatial locations is modestly larger for cross than for iso stimuli. Moreover, the strongest connection in the average connectivity matrix is from excitatory neurons onto VIP neurons within the same subnetwork (Figure 3D). Therefore, modest increases in excitatory drive lead to an increase in VIP activity, causing a decrease in SOM activity, which in turn reduces the inhibition onto excitatory neurons. Thus, excitatory drive is amplified by both recurrent excitation and by the VIP-SOM circuit.

### Feedback drive

In addition to feedforward and local recurrent inputs, feedback inputs may also contribute to contextual modulation. In particular, we have recently shown that excitatory neurons in L2/3, but not L4, have a second receptive field surrounding the ffRF, mediated by feedback excitatory projections from higher visual areas (i.e. feedback receptive field, fbRF; Keller et al., 2020). These feedback projections might provide a source of excitation driving L2/3 excitatory neurons in response to both cross and iso stimuli. VIP but not SOM neurons also appear to have a fbRF (Keller et al., 2020). Since VIP neurons show virtually no responses to iso stimuli (Figure 2G, H), it is unlikely that VIP neurons receive strong feedback projections for such stimuli. We hypothesize that the orientation tuning of the feedback projections targeting VIP neurons may be biased towards the cross orientation, helping VIP to dominate the VIP-SOM circuit for cross stimuli. Further experiments will be necessary to determine the relation of the receptive field properties of the two receptive fields in VIP neurons.

### Conclusions

Contextual modulation represents a fundamental computation to extract meaning from visual scenes. It could support many perceptual phenomena, such as pop-out effects, figure-ground segregation, detection of borders, and object detection (Angelucci et al., 2017; Bergen and Julesz, 1983; Jones et al., 2001; Kapadia et al., 2000; Knierim and van Essen, 1992; Lamme, 1995; Rossi et al., 2001; Schnabel et al., 2018; Seriès et al., 2003; Treisman and Garry Gelade, 1980). Furthermore, the dichotomy between surround suppression and cross-orientation facilitation is consistent with a predictive processing framework (Bastos et al., 2012; Keller and Mrsic-Flogel, 2018), that is, a framework in which the features of a stimulus at a given location can be used to estimate the features of a stimulus at an adjacent location (Rao and Ballard, 1999). Based on natural statistics of the visual environment, the spatial features in a small patch of visual world are likely to be similar to the spatial features in the adjacent patches.

If the stimuli in the surround provide a correct estimate of the stimulus in the center, the response of the neuron can be suppressed, i.e. surround suppression, as there is no need to transmit a signal that is accurately predicted. On the other hand, when the center and the surround differ, the stimuli in the surround provide an incorrect estimate of the stimulus in the center and the signal of the neuron will not be suppressed but passed along, i.e. cross-orientation facilitation. In conclusion, predictive processing is a strong framework for contextual modulation of visual responses in cortical circuits.

## Acknowledgements

We thank M. Mukundan, B. Wong, and L. Bao for technical support, J.I. Glaser for technical advice on the model optimization procedure, and the members of the Scanziani laboratory for helpful discussions of this project. We thank H. Adesnik for the AAV2/9.CAG.Dio.eNpHre3.0.mRuby3.WPRE.SV40 virus and M. Rio for software support. This project was supported by NIH grant U19NS107613 (K.D.M., M.S., M.D., and A.I.), the Howard Hughes Medical Institute (M.S.), the Swiss National Science Foundation grants P300PA_177882 and P2EZP3_162284 to A.J.K and P300PA_177898 to M.M.R, the Gatsby Charitable Foundation (K.D.M. and A.I.), and NSF NeuroNex Award DBI-1707398 (K.D.M. and M.D.). We acknowledge computing resources from Columbia University’s Shared Research Computing Facility project, which is supported by NIH Research Facility Improvement Grant 1G20RR030893-01, and associated funds from the New York State Empire State Development, Division of Science Technology and Innovation (NYSTAR) Contract C090171, both awarded April 15, 2010.

## Author contributions

M.S., A.J.K., and M.S.C. designed the study. A.J.K. and M.M.R. conducted all experiments and experimental data analysis. M.S.C. performed preliminary experiments. M.D. and K.D.M. designed the model. M.D. and A.I. developed the training algorithm of the model. M.D. performed the numerical simulations and analyzed the model results. A.J.K., M.M.R., M.D., K.D.M., and M.S. wrote the manuscript.

## Methods

### Animals

All experimental procedures were conducted in accordance with the regulation of the Institutional Animal Care and Use Committee of the University of California, San Francisco. The mice were housed on a reverse light cycle (light/dark cycle: 12/12 hrs). At the start of the experiments, all mice were older than 2 months. Mice were of either sex and were of the following genotype:

Gad2-IRES-cre (GAD2^tm2(cre)Zjh^; JAX:010802) × Ai14 (Gt(ROSA)26Sor^tm14(CAG-tdTomato)Hze^; JAX:007914) for imaging of layer 2/3 (L2/3) excitatory neurons (9 mice; Figures 1, 3, S1, S2); Scnn1a-Tg3-cre (Tg(Scnn1a-cre)3Aibs/J; JAX:009613) and Scnn1a-Tg3-cre (Tg(Scnn1a-cre)3Aibs/J; JAX:009613) × Ai148 (Igs7^tm148.1(tetO-GCaMP6f-CAG-tTA2)Hze^; JAX:030328) for imaging layer 4 (L4) excitatory neurons (4 mice and 1 mouse, respectively; Figures 1, 3, S2); Sst-IRES-cre (Sst^tm2.1(cre)Zjh^; JAX:028864) × Ai14 (Gt(ROSA)26Sor^tm14(CAG-tdTomato)Hze^; JAX:007914) for imaging of L2/3 somatostatin-expressing neurons (SOM; 13 mice; Figures 2, 3, S2); PV-cre (Pvalb^tm1(cre)Arbr^; JAX:017320) × Ai14 (Gt(ROSA)26Sor^tm14(CAG-tdTomato)Hze^; JAX:007914) for imaging of L2/3 parvalbumin-expressing inhibitory neurons (PV; 10 mice; Figures 2, 3, S2); VIP-IRES-cre (Vip^tm1(cre)Zjh^; JAX:010908) × Ai14 (Gt(ROSA)26Sor^tm14(CAG-tdTomato)Hze^; JAX:007914) for imaging of L2/3 vasoactive-intestinal-peptide-expressing inhibitory neurons (VIP; 7 mice; Figures 2, 3, S2); VIP-IRES-cre (Vip^tm1(cre)Zjh^; JAX:010908) for optogenetic manipulation of VIP neurons and imaging putative excitatory and VIP neurons (8 mice; Figures 4, S4); and VIP-IRES-cre (Vip^tm1(cre)Zjh^; JAX:010908) × Sst-IRES-Flp (Sst^tm3.1(flpo)Zjh^; JAX:028579) for optogenetic manipulation of VIP neurons and imaging SOM neurons (8 mice; Figure 4).

### Viruses

Viruses were typically diluted to use titers of approximately 5 × 10^12^ genome copies/ml and 50 nl were injected at each injection site (3 to 5 sites per mouse) and each depth (2 from 350 to 200 μm below the pial surface). We injected the following viruses:

AAV2/1.ef1a.GCaMP6f.WPRE (FMI Vector Core Facility), AAV2/1.ef1a.DIO.GCaMP6f.WPRE (FMI Vector Core Facility), AAV2/1.CAG.CGaMP6f (Janelia Vector Core), AAV2/9.syn.GCaMP7f (Addgene), AAV2/1.ef1a.fDIO.GCaMP6s (Janelia Vector Core), AAV2/5.CBA.Flex.ArchT-tdTomato.WPRE.SV40 (University of Pennsylvania Vector Core), AAV2/1.CAG.Flex.rc[Jaws-KGC-GFP-ER2] (Janelia Vector Core), AAV2/9.CAG.Dio.eNpHre3.0.mRuby3.WPRE.SV40 (H. Adesnik), and AAV2/9.ef1a.F-Flex.tdTomato (Xue et al., 2014).

### Surgery

Mice were anesthetized with 2% isoflurane or with a mixture of Fentanyl (West-Ward Pharmaceuticals, 0.05 mg/kg), Midazolam (Akorn, 5.0 mg/kg) and Dexmedetomidine (Zoetis, 0.5 mg/kg), injected subcutaneously. Mice’s body temperature was monitored and kept constant. To prevent the eyes from drying, a layer of lubricant ointment (Rugby) was applied. The skin above the skull was disinfected with povidone iodine. A craniotomy was made over the right visual cortex (3 to 4.5 mm in diameter) and viruses were injected with a micropump (UMP-3, World Precision Instruments) at a rate of 2 nl/s. The craniotomy was then sealed with a glass coverslip using cyanoacrylate glue and a headplate was attached. To reverse the anesthesia induced by the Fentanyl-Midazolam-Dexmedetomidine mixture, a mixture of Naloxone (Hospira, 1.2 mg/kg), Flumazenil (West-Ward Pharmaceuticals, 0.5 mg/kg), and Atipamezol (Zoetis, 2.5 mg/kg) was injected subcutaneously after the surgical procedures.

### Visual stimulation

Visual stimuli were generated using the open-source Psychophysics Toolbox based on Matlab (MathWorks). Stimuli were presented at 15 cm to the left eye on a gamma-corrected LED-backlit LCD monitor (DELL) with a mean luminance of 20 cd/m^2^. For experiments using a resonant scanner, the power source of the monitor’s LED backlight was synchronized to the resonant scanner turnaround points (when data were not acquired) to minimize light leak from the monitor (Leinweber et al., 2014). We presented drifting sinusoidal gratings (2 Hz, 0.04 cycles/°, 100% contrast) unless stated otherwise. The trial structure of all stimulus sessions (receptive field mapping, orientation tuning, *et cetera*) was block randomized (the block size was given by the total number of parameter combinations).

#### Receptive field mapping

Stimuli consisted of a circular grating patch on a gray background (typically set to 20° in diameter). Stimuli were presented for 1 s at a single direction or for 2 s at the 4 cardinal directions (0.5 s each). Stimulation periods were interleaved by 2 s of gray screen. We recorded 5 to 10 trials per stimulus condition.

#### Orientation tuning

We presented gratings of at least 10° diameter drifting in 8 directions (5 to 10 trials). Stimulus time was 2 s interleaved with 4 s of gray screen.

#### Size tuning

Patches of gratings were displayed at up to 9 different sizes, linearly spaced from 5° up to 85° in diameter (10 trials per size) centered on the classical feedforward receptive field (ffRF). Stimulation time was 2 s interleaved by 4 s of gray screen. Stimuli were either presented at a single direction or at the 4 cardinal directions (0.5 s each).

#### Contextual modulation

We presented patches of gratings (10° to 30° in diameter) on a gray background (center stimulus), full-field gratings (iso stimulus), and patches of gratings (10° to 30° in diameter) on cross-oriented fullfield gratings (cross stimulus). Stimulation time was 2 s interleaved by 4 s of gray screen. Trials with optogenetic stimulation had an additional 1 s pre-stimulus and post-stimulus gray screen during which the optogenetic light source was turned on and the total number of trials was doubled (**Optogenetics** below).

### Behavioral monitoring

All mice were habituated (3 to 5 days) to the experimental setup before starting experiments. During all experiments, we recorded the positions of the left eye using a CMOS camera (DMK23UM021, Imaging Source) with a 50 mm lens (M5018-MP, Moritex), tracked the running speed of the mouse, and monitored its general behavior using a webcam (LifeCam Cinema 720p HD, Microsoft).

### Two-photon calcium imaging

Imaging was performed using either a galvanometric-scanner based MOM (Sutter) or a resonant-scanner based (8 kHz) Bergamo II two-photon microscope (Thorlabs), both controlled by ScanImage (Vidrio). Using the MOM system, we acquired images of 128 × 128 pixels at a single depth at 5.92 Hz frame rate. With the Bergamo II, we acquired images of 380 × 512 pixels at 1 or 4 depths at 40 Hz or 8 Hz frame rate, respectively. We obtained similar results with both systems, so all data were pooled. The illumination light source was a Ti:sapphire laser (Chameleon Ultra II, Coherent) used at an excitation wavelength of 910 nm. The laser power under the objective (16×, Nikon) was typically set to 30 mW and never exceeded 50 mW (laser pulse width 140 fs at a repetition rate of 80 MHz).

### Optogenetics

To silence VIP neurons, we used a 594 nm laser (OBIS 594 LS 100 mW, Coherent). We modified the Bergamo II microscope (Thorlabs) to combine optogenetic manipulation with two-photon calcium imaging. A lens (LA1805-B, Thorlabs) was placed in the optogenetic stimulation light path to defocus the light at the imaging plane. We used a dichroic mirror (DMBP740B, Thorlabs) to combine two-photon laser and optogenetic stimulation light. Moreover, we used a second dichroic mirror (FF555-Di03-25×36, Semrock) to split the green fluorescent protein (GFP) emission from both the two-photon and optogenetic light sources. The laser for optogenetic stimulation was synchronized to the resonant scanner turnaround points (when data were not acquired) to minimize light leak from the monitor (Attinger et al., 2017; see **Visual Stimulation** for timing within a trial). The 594 nm laser power under the objective did not exceed 18 mW.

### Data analysis

All data were analyzed using custom-written code in Matlab (MathWorks).

#### Two-photon calcium imaging

We analyzed two-photon calcium imaging data as described previously (Keller et al., 2020). Briefly, data were full-frame registered using custom-written software (https://sourceforge.net/projects/iris-scanning/). We selected the neurons semi manually, based on mean and maximum projection images. We calculated the raw fluorescence traces as the average fluorescence of all pixels within a selected region of interest for each frame. Fluorescence changes (ΔF/F) were calculated as described elsewhere (Dombeck et al., 2007). All stimulus evoked responses were baseline subtracted (1 s pre-stimulus interval).

#### Response amplitude

The response amplitude to a stimulus was computed as the average response over the duration of the stimulus presentation (excluding the first 0.5 s of each trial due to the delay and slow rise of calcium indicators). We defined significant responses as responses that exceeded a z-score of 3.29 (corresponding to p < 10^-3^) or 5.33 (corresponding to p < 10^-7^; for experiments in L4).

#### Receptive field mapping

To estimate the center of the receptive field, we fitted the responses to patches of gratings with a two-dimensional Gaussian. We excluded neurons if they failed to have at least one significant trialaveraged response within 10° of their estimated ffRF centers. Additionally, except for the ‘surround group’ (see **Computational model**), we excluded neurons if their estimated ffRF centers were not within 10° of the stimulus centers of the stimuli used for estimating size tuning, orientation tuning, *et cetera.* Neurons of the ‘surround group’ had estimated receptive field centers that were at least 15° away from the centers of the stimuli.

#### Size tuning

We fitted the integral over a difference of Gaussians. This fit was used to estimate the neurons’ preferred sizes. We approximated the ffRF size by the size of the patch of gratings evoking the largest response (size tuning fits were bound to the interval 0.1 to 90.1°).

#### Orientation tuning

We fitted a circular sum of Gaussians with a peak offset of 180° and equal tuning width (full width at half maximum of the Gaussian fit). When the preferred orientations of neurons were relevant, we excluded neurons with an R^2^ goodness-of-fit of 0.3 or below.

#### Contextual modulation

To estimate the contextual modulation of excitatory, VIP, SOM, and PV neurons, we used a center patch diameter of 20°. We calculated a contextual modulation index defined as the difference between the activity to cross and iso stimuli divided by the sum of the two. To estimate the effect of silencing VIP neurons on the contextual modulation of putative excitatory neurons, neurons were only considered if their preferred size was within 10° of the center-patch diameter. Note that for these experiments, the center-patch diameter was set to a size between 10° and 30°. Population-averaged responses to center, iso and cross stimuli were calculated based on normalized responses (Figures 3C, S2A). To this end, trial-averaged responses of every neuron were normalized by the maximum responses to center, iso, cross, and receptive field mapping stimuli.

#### Surround suppression

Surround suppression was computed as one minus the responses to iso (or cross) divided by the responses to center stimuli. Neurons with a negative response to center were excluded from this analysis.

#### Baseline

We estimated the baseline activity as the difference between the average fluorescence change during baseline periods (averaged over all 1 s pre-stimulus intervals) and the lower quartile of the overall trace of fluorescence changes. To compute the population-averaged baseline activity, we excluded neurons with an estimated baseline activity of more than 3 standard deviations above the median.

### Computational model

We developed a model reproducing the responses of the 5 different neuronal types from which we recorded (L2/3 excitatory, VIP, SOM, and PV inhibitory neurons and L4 excitatory neurons). Each unit of the circuit represented the average activity of a given neuron type, integrated in a ‘subnetwork’ with the other unit types. Four such subnetworks were each assigned to one of two spatial locations (each considered the ‘surround’ of the other) and one of two preferred orientations (orthogonal to each other). We consequently obtained a total of 20 units, 5 unit types in 4 subnetworks. We optimized the synaptic strengths between these model units to match their responses to those observed experimentally. We obtained many solutions by using many sets of pseudo data (obtained by perturbing the experimental data by random noise with standard deviation proportional to the measured error).

#### Experimental data

To model the average activity of our 20 units split across the 4 subnetworks, we divided our experimental data set into 4 subgroups: ‘Centered and preferred orientation’, ‘centered and orthogonal orientation’, ‘surround and preferred orientation’, and ‘surround and orthogonal orientation’ (for details, see **Data analysis**). The population-averaged responses of the 5 neuron types within each subgroup were the targets of the corresponding 5 units within the 4 subnetworks of our model. Centered neurons were those with ffRFs aligned with the location of the center stimulus, i.e. ffRFs no more than 10° from the stimulus center. Surround neurons were those with ffRF offset from the location of the center stimulus, i.e. ffRF centers were at least 15° from the stimulus center. ‘Preferred orientation’ neurons were those with preferred orientation within 45° of the center stimulus orientation. ‘Orthogonal orientation’ neurons were those with preferred orientation more than 45° from the center stimulus orientation. Population-averaged responses of the neurons of each of the 5 types within each of the 4 subgroups were obtained for each of 4 stimulus conditions (spontaneous activity or presentation of the center, iso, or cross stimuli). Hence, the goal of the model was to fit model responses to the matrix of these experimentally observed mean responses, 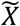 whose elements 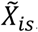, where *i* corresponded to one of the 20 units and s corresponded to one of the 4 stimulus conditions (Figure S2A).

#### Model parameters

Connections between L4 excitatory units and the other model units were unidirectional, as L4 was considered as an input to the subnetwork. The L4 unit of a given subnetwork was restricted to project only to excitatory (Exc) and PV units of the same spatial location, but of either preferred orientation (Adesnik et al., 2012; Karnani et al., 2016). Within a subnetwork, there were 16 possible recurrent connections between L2/3 excitatory and inhibitory units, of which we disallowed 5 that were deemed negligible based on electrophysiological measurements (Pfeffer et al., 2013). We disallowed the following connections: VIP → Exc, VIP → PV, VIP → VIP, SOM → SOM, and PV → SOM. Thus, each subnetwork received 11 recurrent connections and two L4 connections from within its own subnetwork, a total of 13 connections per subnetwork (W^(1)^ in Figure 3D). The same set of connections was also allowed from the opposite orientation at the same location (W^(2)^ in Figure 3D), making 26 connections to a given subnetwork from its own spatial location. Projections across spatial locations were only allowed from L2/3 excitatory units to all four L2/3 unit types, adding 8 additional connections received by each subnetwork (W^(3)^ and in Figure 3D; the connections from inhibitory and L4 neurons were all set to zero and therefore not displayed in Figure 3D). In total, we thus allowed 34 non-zero connections per subnetwork.

The overall 16 × 20 weight matrix was composed of the 4 × 5 submatrices in the following convolutional structure

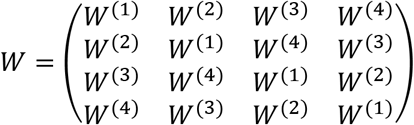

This structure meant that each subnetwork can be considered as the surround of the other and each orientation as the orthogonal of the other. This symmetry across domains allowed us to keep the total number of parameters at 34.

The above matrix *W* was defined in a basis in which the 20 rates were arranged as (Exc, PV, SOM, VIP, L4) of network 1, then network 2, then 3, then 4. We rearranged these weights and rates, letting *A* be the 16 × 16 matrix of recurrent weights between the sixteen L2/3 units, found from *W* by keeping only the first 4 columns of each of the *W*^(*i*)^; and *B* be the 16 × 4 matrix of projections from L4 units to the sixteen L2/3 units, found from *W* by keeping only the last column of each of the *W*^(*i*)^. Then, in this rearranged basis, *W* became *(A, B),* and acted on a rate vector 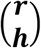 whose first 16 elements ***r*** were the rates of the L2/3 units and whose last 4 elements ***h*** were the rates of the L4 units (we use bold font to indicate vectors, and capital letters to indicate matrices).

#### Rate equations

The rate equations for the units in the network for a particular stimulus s were

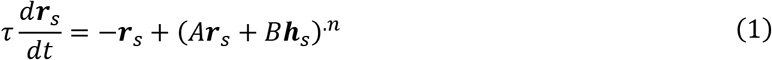

where the element-wise operation 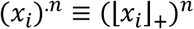 corresponded to the input-output function, a rectified power law with exponent *n* = 2 (Ahmadian et al., 2013). The 16-vector ***r**_s_* specified the activities of the L2/3 units to stimulus *s,* while the 4-vector ***h**_s_* specified the activities of the L4 units to that stimulus. The time constant was set to *τ* = 10 ms. We denoted the combination of L2/3 and L4 units by 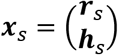 while, again, the combination of recurrent and feed-forward weights is *W* = (*A, B*). We used *X, R*, and *H* to refer to the matrices whose columns are the vectors ***x**_s_*, ***r**_s_*, or ***h**_s_*, respectively, across all stimuli *s*.

#### Cost function

For each stimulus *s*, we denoted the experimentally measured mean responses as 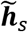 for L4 and 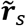 for L2/3. Our model found inputs (L4 responses) 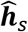 and synaptic weights that produced a steady-state response denoted by 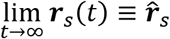. The cost function of the model demanded that the inputs and responses should have minimal summed-weighted-squared error relative to the experimental measurements, subject to certain regularization terms:

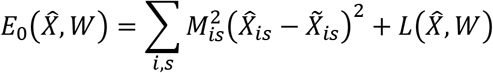

Here, *M_is_* was a weight matrix that represented our uncertainty over the responses. More specifically *M_is_* = *σ_0_β_i_/σ_is_*, where *σ_is_* was the standard error of the responses 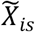 measured experimentally, *β* was a multiplicative factor to weight errors in certain unit types more than others, and *σ*_0_ = 〈*σ_is_/β_j_*〉_*is*_ was a normalization factor, where 〈*z_is_*〉_*is*_ indicated an average of *z_is_* over *i* and *s*. We chose *β_i_* = 1 for L2/3 excitatory, PV and VIP neurons, *β_i_* = 2.5 for SOM and *β_i_* = 5 for L4 excitatory neurons. We used larger *β_i_* for units that we found harder to fit. Intuitively, this fitting difficulty might arise from the fact that L4 and SOM neurons had the most distinct response patterns compared to other neuron types. *L* represented the sum of all regularization terms, defined as:

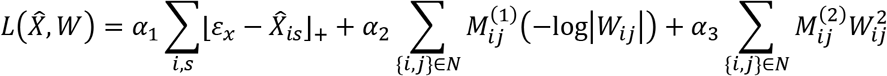

The first regularization factor, using *α*_1_ = 0.02, nudged the responses 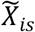 above a minimal threshold *ε_x_* = 0.01, since 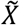 corresponded to estimated firing rates and were thus non-negative. The second and third factors were applied only to the 34 weights that were allowed to be nonzero, as specified above; this set of weights was (1) designated by *N.* The second factor, using *α*_2_ = 0.05, nudged weights with a corresponding positive value of 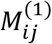 to prevent them from being too close to zero. The elements 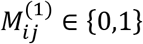 were non-zero for almost all allowed connections between units sharing orientation preference: all of those from the same spatial location of the ffRF (*W*^(1)^), and those from the surround to excitatory units and to SOM units (two of the 4 non-zero elements of *W*^(3)^; among the potential targets of projections across spatial locations, we only pushed L2/3 excitatory and SOM units away from zero because those have well-established evidence for substantial projections across spatial locations; Adesnik et al., 2012). The third factor, starting with *α*_3_ = 0.01, nudged weights with a corresponding positive value of 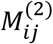 towards zero. The elements 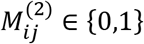 were non-zero for all allowed connections between units having different preferred orientations (*W*^(2)^ and *W*^(4)^). In other words, we wanted to discourage strong cross-orientation connections and encourage strong connections among those allowed in *W*^(1)^ and *W*^(3)^.

#### Training a recurrent neural network with custom weight initialization

We noted that equation (1) corresponded to a recurrent neural network (RNN). This allowed us to train the RNN to find the best solutions, i.e. the weights *W_ij_* and the inputs *H_ij_*, using backpropagation through time (BPTT; Pascanu et al., 2013; Rumelhart et al., 1986). The training of a neural network was highly sensitive to its initial weights (He et al., 2015) and in general we observed that starting from random initial conditions would often lead to unstable solutions. This might stem from the fact that RNN training was prone to gradient vanishing and gradient explosion (Bengio et al., 1994), especially for a large number of time steps. As a first step, we therefore found stable solutions which would approximately match the data using non-negative least square (NNLS) regression, which we used as initial conditions of the BPTT. In a previous study (Dipoppa et al., 2018), we used a NNLS to infer the optimal synaptic strengths of a model evolving a dynamical equation similar to equation (1) such that the model would match the experimental data. Here we similarly inferred optimal strengths for matching the model to pseudo data 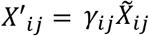, randomly generated using the random matrix 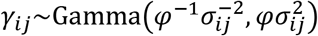 with *φ* = 5, such that 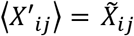 and 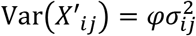. We solved the convex problem of minimizing the following cost function, *E*_1_, which made *X*’ as close as possible to a fixed point of equation 1 subject to regularization, as an approximation of minimizing *E*_0_(*X*’, *W*):

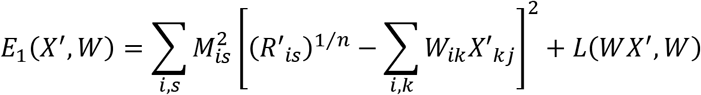

Here *R*’ was the *R* component of 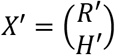. We generated *N_NNLS_* = 2,500,000 different sets of pseudo data {*X*’}. We then used the trust region reflective algorithm to solve the problem 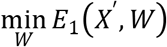 starting from initial conditions *W_ij_*~Gamma(1,1) and with boundaries 0 < |*W_ij_*| < 10. After obtaining a set of optimal parameters {*W_NNLS_*} for each set of pseudo data, we let the system evolve following equation (1) and obtained the fixed points (if they existed), discarding all solutions that had at least one of the 20 rates >10 or < *ε_x_*. This produced the set of fixed points 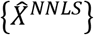. Note that the 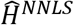 portion of 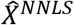 was unchanged from its original perturbed value 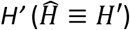. We then recomputed the error 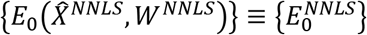.

We selected the 50,000 best solutions 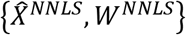 sorted by 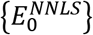 as starting parameters for the BPTT. We defined the following cost function for the BPTT:

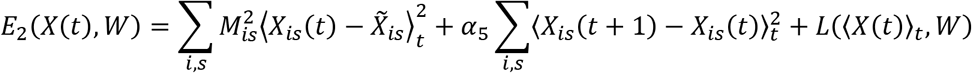

where *X_is_*(*t*) corresponded to the dynamics of the system at each time step *t* and where the average over *t*, 〈·〉_*t*_, was computed over the last *T* = 200 time steps of the dynamics. The factor with *α*_5_ = 1 punished large values of the derivative of *X* to ensure that the system reached a fixed point. Independently of the stimulus condition, for each run of the dynamics (termed an ‘epoch’), the starting point was 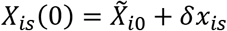, where *s* = 0 corresponded to the spontaneous activity and 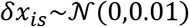 was the random perturbation following a Gaussian distribution. We used time steps of Δ*t* = 2 ms. An epoch consisted of evolving equation (1) using the Euler scheme:

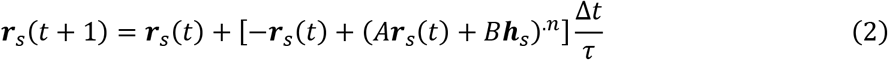

for 500 time-steps. To compute the gradient of the loss equation *E*_2_(*X*(*t*), *W*) over *W* and the *H* portion of *X* of the discretized system given by equation (2) over the last *T* = 200 time steps of the dynamics, we used automatic differentiation methods provided by the pytorch library in Python. Optimization was carried out by the ADAM optimizer (Kingma and Ba, 2017). To improve convergence to a solution, we employed a triangular learning rate policy (Smith, 2017) at a base learning rate of 3 × 10^-4^, a maximum learning rate of 3 × 10^-3^, 100 training epochs for the increasing part of the cycle, 200 training epochs for the decreasing part of the cycle. We also used a patience parameter of 1000 epochs. If the error did not improve over this length of time, the training procedure of the BPTT would stop. If not interrupted, the model was trained for 10,000 epochs. After running the BPTT for the best 50,000 starting conditions of the NNLS 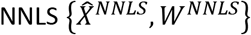, we obtained a new set of inferred weights and rates 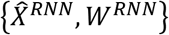. Note that in contrast to the NNLS, the 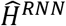 portion of 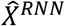 was learned. Of these, we selected the top 15 or top 100 solutions sorted by the smallest error 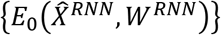 for further analysis.

### Analysis of the computational model

#### Comparison with the data

The model was trained to reproduce the estimated firing rate of the units *X* (Figure S2A). For better comparison to the experimental data, we showed the baseline subtracted neuronal responses Δ*X_is_* = *X_is_* — *X*_*i*0_, i.e. with the spontaneous activity s *=* 0 subtracted (Figure 3C). Similarly, we used Δ*X_is_* to compute CMIs and the difference between control and optogenetic conditions (Figure 3F, Figure S3). Using Δ*X_is_* instead of the neuronal activity *X_is_* did not appreciably change the results.

#### Model of surround modulation index

We observed a large variability of Δ*CMI* (the change in contextual modulation index upon VIP silencing) of L2/3 excitatory units across the top *N_s_* = 100 solutions, with the majority of the top 15 solutions having a positive value of Δ*CMI* (Figure S3B, C). We asked whether we could predict this variability from the difference in connection weights across these solutions. We defined ***q**_p_* as the vectorized form of the unique (not repeated by the convolution) and non-zero connection weights ({*i,j*} ∈ *N*, see **Computational model**) of the *p^th^* solution 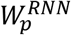, and Δ*CMI_p_* as the change in contextual modulation index of centered and preferred L2/3 excitatory units for the *p^th^* solution. We then ran *N_r_* = 100 lasso (least absolute shrinkage and selection operator) regressions with 10-fold cross validation (i.e. *L*_1_-regularized linear regression), each one with a different random seed, relating the predictors in ***q**_p_* to the responses in Δ*CMI_p_*. Across the *N_r_* regressions Δ***CMI*** = *β*_0_+*β*_1_*Q*, where Δ***CMI*** = (Δ*CMI*_1_, …, Δ*CMI_N_s__*) and *Q* = (***q***_1_, …, ***q**_N_s__*), we selected the sparsest solution, i.e. the one with the lowest number of non-zero coefficients *β*_1_ (Figure S3E).

### Statistics

We used two-sided Wilcoxon rank-sum tests for independent group comparisons, and two-sided Wilcoxon signed-rank tests for paired tests and single group analysis. No statistical methods were used to predetermine experimental sample sizes.

**Figure S1.**
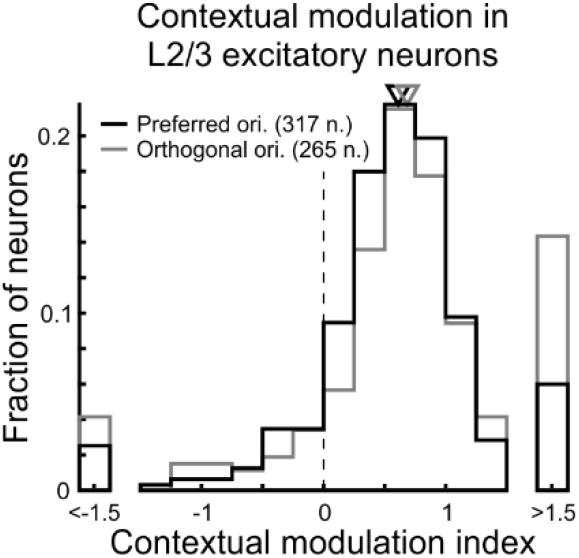
Contextual modulation in L2/3 excitatory neurons separated by orientation preference. Related to Figure 1. Iso stimuli and the center of cross-surround stimuli were both presented at the same orientation. Neurons were split in two groups based on their orientation tuning, one group with neurons having a preferred orientation similar to that of the presented orientation and another group with neurons having a preferred orientation orthogonal to that of the presented orientation (see Methods). Contextual modulation index distributions for preferred orientation (black) and for orthogonal orientation (gray). Triangles above histograms indicate median. Single-distribution two-sided Wilcoxon sign-rank test; preferred orientation: p < 10^-10^; orthogonal orientation: p < 10^-10^; 317 and 265 neurons in 9 mice, respectively.

**Figure S2.**
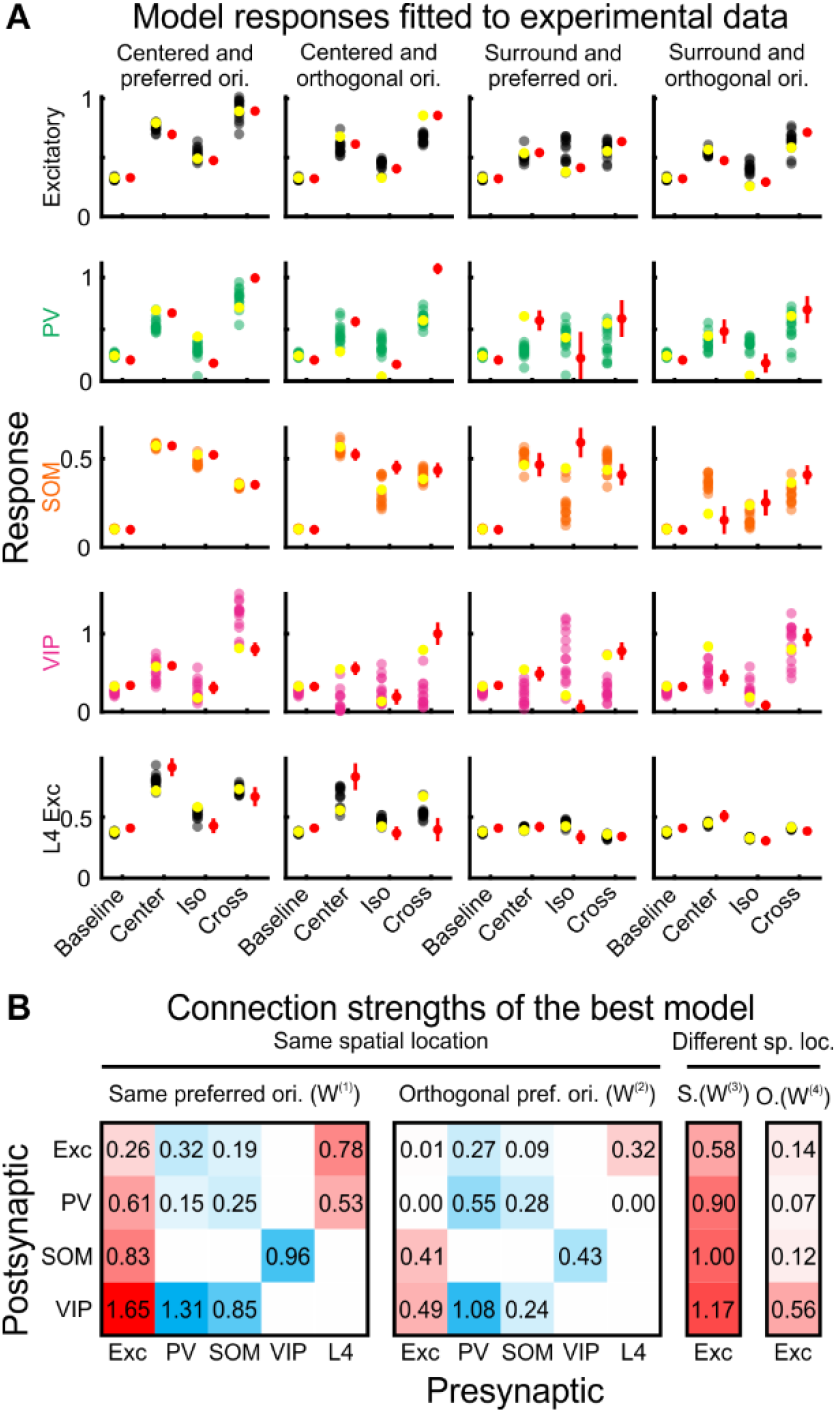
Model response fits and example model. Related to Figure 3. (A) Spontaneous activity, and responses to center, iso, and cross stimuli of the 5 unit-types for the top 15 models (not baseline subtracted; see Methods). Units were fit to experimental data considering two spatial locations and two preferred orientations (columns). ‘Centered’: Neuron ffRFs aligned with the stimulus location. ‘Surround’: Neuron ffRFs offset from the stimulus location. ‘Preferred orientation’: Neurons with preferred orientation matching the stimulus orientation. ‘Orthogonal orientation’: Neurons with preferred orientation orthogonal to the stimulus orientation. Yellow symbols represent example model in (B). Each dot represents the activity of a unit from a single model. Red symbols represent experimental data (mean ± SEM; 911, 317, 265, 180, and 172 L2/3 excitatory neurons in 9 mice for baseline and the 4 functional groups (columns), respectively; 80, 48, 21, 7, and 8 PV neurons in 10 mice; 303, 200, 60, 24, and 22 SOM neurons in 13 mice; 74, 30, 10, 20, and 16 VIP neurons in 7 mice; 96, 22, 13, 29, and 34 L4 excitatory neurons in 5 mice). (B) Connection strengths of the best model. Excitatory connections are represented in red, inhibitory connection in blue. The 4 matrices correspond to W^(1)^, W^(2)^, W^(3)^, and W^(4)^ in Figure 3B.

**Figure S3.**
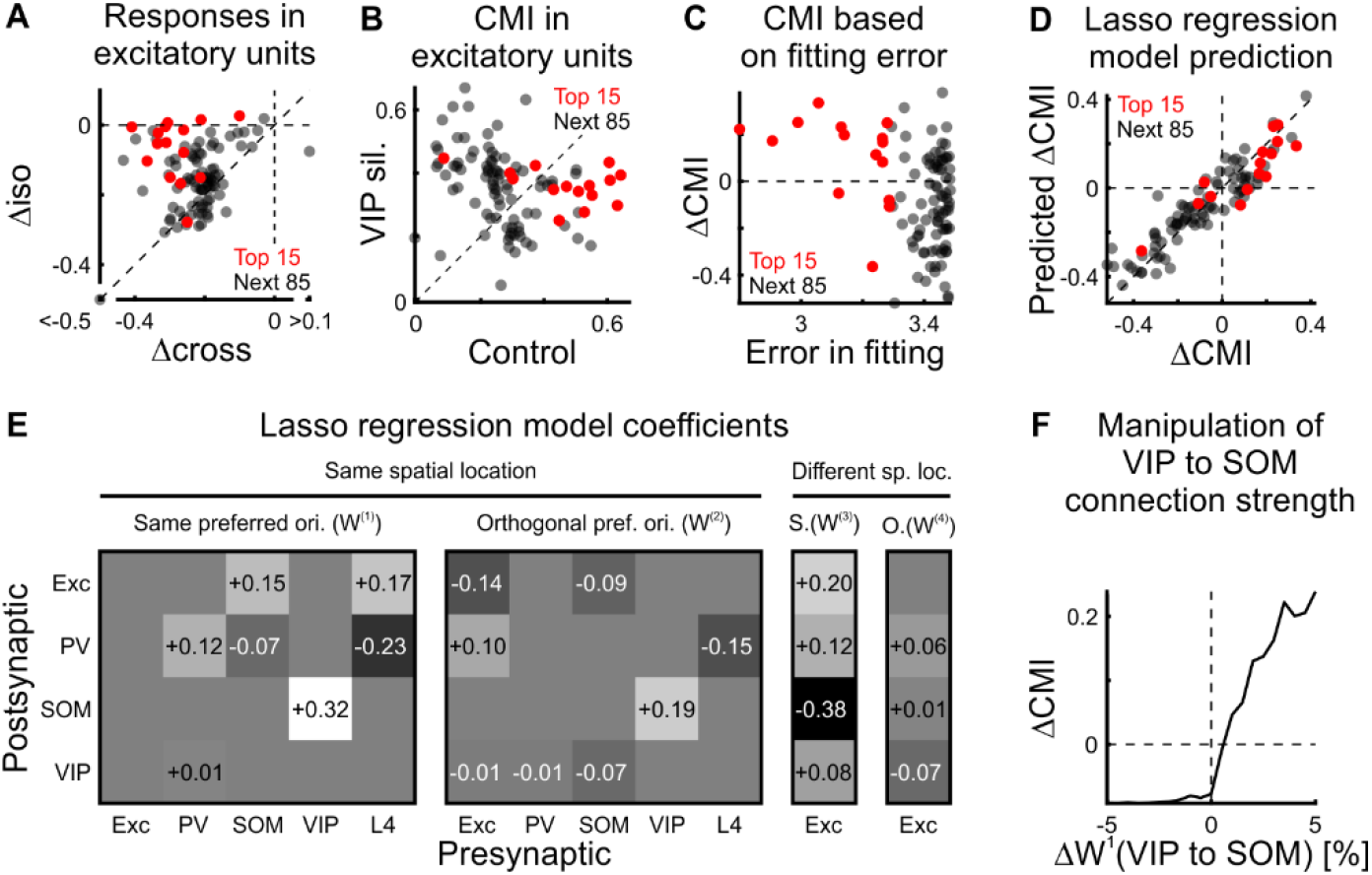
Analysis of top 15 and next 85 recurrent neural network models. Related to Figure 3. (A-D) Responses of excitatory units to stimuli centered on the spatial location of their ffRF and presented at their preferred orientation for the top 15 models (red) and the next 85 models (black). (A) Changes in responses to iso and cross stimuli in excitatory units upon VIP silencing. Paired two-sided Wilcoxon sign-rank test; top 15 models, p = 1.2 × 10^-4^; next 85 models, p < 10^-6^. (B) Changes in CMI in excitatory units upon VIP silencing. Paired two-sided Wilcoxon sign-rank test; top 15 models, p = 0.030; next 85 models, p < 10^-4^. (C) Change in CMI upon VIP silencing (ΔCMI) plotted against error in fitting (see Methods). (D) Lasso (least absolute shrinkage and selection operator) regression model prediction of ΔCMI. The prediction of ΔCMI for each recurrent neural network (RNN) model solution is given by the sum of the lasso regression model coefficients, represented in (E), times the deviation of the RNN model from the average connection strength of the top 100 RNN models. (E) Lasso regression model coefficients. (F) Median ΔCMI of the top 100 RNN models increases if the VIP to SOM connection in W^(1)^ is strengthened (same spatial location and same preferred orientation).

**Figure S4.**
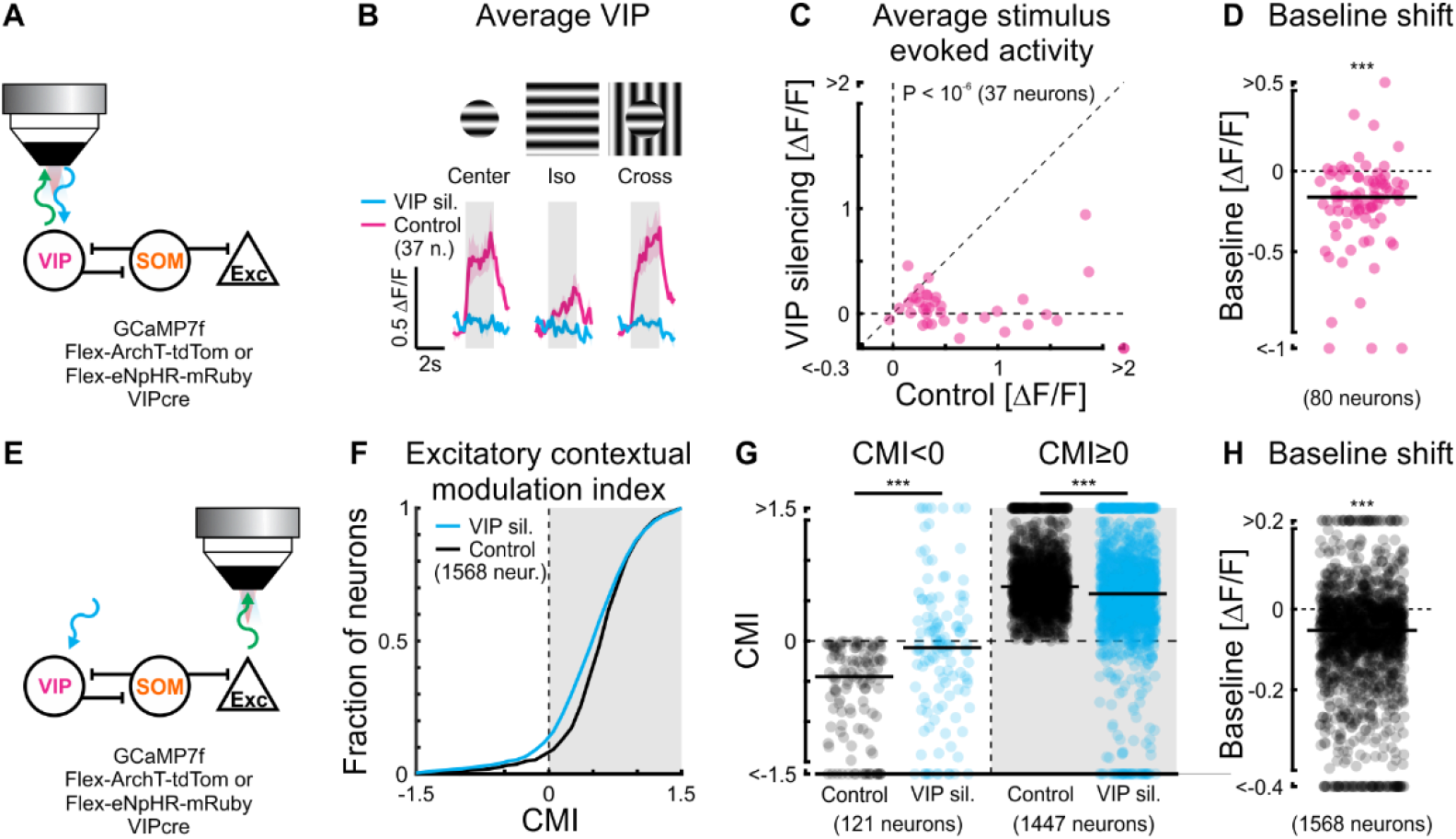
Silencing VIP neurons and its effect on excitatory neuron responses to 100% contrast stimuli. Related to Figure 4. (A) Experimental setup. We conditionally expressed an inhibitory opsin, ArchT or eNpHR, in VIP neurons and unconditionally expressed a calcium indicator, GCaMP7f. (B) Population-averaged calcium responses of VIP neurons with and without silencing VIP neurons (37 neurons in 6 mice). Here, stimuli were presented at 50% contrast. (C) Scatter plot of stimulus-averaged responses (center, iso and cross at 50% contrast) in VIP neurons with and without silencing VIP neurons. Paired two-sided Wilcoxon sign-rank test; p < 10^-6^; same neurons as in (B). (D) Baseline shift in VIP neurons upon silencing VIP neurons. Single-distribution two-sided Wilcoxon sign-rank test; ***: p < 10^-9^; 80 neurons in 8 mice. (E) Same experimental setup as in (A). (F) Cumulative sum of CMIs in putative L2/3 excitatory neurons (1568 neurons in 8 mice). Here, stimuli were presented at 100% contrast. (G) Upon silencing VIP neurons, putative L2/3 excitatory neurons with a negative CMI increased their CMI and those with a positive CMI decreased their CMI (gray shading). Paired two-sided Wilcoxon sign-rank; CMI < 0 and CMI ≥ 0, ***: p < 10^-10^; 121 and 1447 neurons, respectively, in 8 mice. (H) Baseline shift in putative L2/3 excitatory neurons upon silencing VIP neurons. Single-distribution two-sided Wilcoxon sign-rank test; ***: p < 10^-10^; same neurons as in (F).

